# Opportunities for improved surveillance and control of infectious diseases from age-specific case data

**DOI:** 10.1101/545319

**Authors:** Isabel Rodríguez-Barraquer, Henrik Salje, Derek AT Cummings

## Abstract

One of the challenges faced by global disease surveillance efforts is the lack of comparability across systems. Reporting commonly focuses on overall incidence, despite differences in surveillance quality between and within countries. For most immunizing infections, the age-distribution of incident cases provides a more robust picture of trends in transmission. We present a framework to estimate transmission intensity for dengue virus from age-specific incidence data, and apply it to 363 administrative units in Thailand, Colombia, Brazil and Mexico. Our estimates correlate well with those derived from seroprevalence data (the gold-standard), capture the expected spatial heterogeneity in risk, and correlate with known environmental drivers of transmission. We show how this approach could be used to guide the implementation of control strategies such as vaccination. Since age-specific counts are routinely collected by many surveillance systems, they represent a unique opportunity to further our understanding of disease burden and risk for many diseases.

## Introduction

A fundamental challenge of disease surveillance systems is how to transform data that is routinely collected into useful, actionable evidence that can inform control interventions. Disease surveillance systems typically focus on analyzing aggregate counts of cases over defined time periods to stratify the risk of populations(1). However, the use of raw case counts, or incidences, as a measure of disease risk can frequently be misleading because the quality of surveillance often differs significantly both across countries and within regions of a country making comparisons inappropriate. Areas with more complete reporting of disease may inaccurately appear to have more disease simply because reported cases scale linearly with completeness of reporting. Surveillance systems may change over time (e.g. improving in completeness over time) making comparisons difficult or even impossible. These problems are particularly troubling when examining disease trends or ranking regions according to their disease risk. Recent efforts have tried to improve quantification of disease burden by pooling numerous sources of data. For example, disease mapping methods that combine disease presence/absence data, environmental covariates and available incidence data (from cohort or cross-sectional studies) have been used to predict spatial limits and global case counts for diseases including malaria, dengue and several neglected tropical diseases. (2-5) However, while these methods are successful in identifying boundaries of endemic areas, the robustness of these approaches to quantify transmission within endemic areas hasn’t been validated.

For most immunizing diseases, serological surveys of immunity are regarded as the gold standard to measure the susceptible fraction and infer the extent of transmission, as they provide a direct measure of the proportion of the population that has been infected at a single point in time or between two or more time points. This is particularly useful for diseases such as dengue, influenza or Zika, where the asymptomatic to symptomatic ratios are large or unknown. Methods to estimate transmission parameters from age-stratified serological data have been available for many years(6) and have been used to analyze trends in transmission for multiple diseases including measles(7), hepatitis A(8), dengue(9-12), pertussis(13), influenza(14), malaria(15) and chikungunya(16). While well conducted age-stratified serological surveys can provide an unbiased measure of the susceptibility profile of a population and transmission parameters, they are generally not part of routine surveillance activities and are therefore only available for a limited number of locations and time points. For example, an extensive review of the dengue literature published recently found only three population-based serosurveys conducted in Brazil over the past 10 years, despite being the country that currently reports the largest number of cases worldwide(17). Similarly, only one and three studies were identified for the Philippines and Thailand, respectively, even though the burden of dengue is very high(17). Thus, the picture provided by serology is very incomplete and limited when trying to characterize and compare local and global trends.

While aggregated case-counts can be misleading when quantifying disease risk, the age-distribution of incident cases contains a lot of information on the age-specific susceptibility of the population. Importantly, the age-distribution of cases is also largely robust to under-reporting, facilitating the comparison between locations or over time. By combining age-specific incidence data and mechanistic models of how population immunity is acquired over time, it is possible to estimate key transmission parameters and obtain a much more accurate picture of the local and global burdens of disease. Since age-specific counts are routinely collected by surveillance systems as part of standard practice, they represent a missed opportunity to further our understanding of epidemic patterns for many diseases.

Here, we use dengue virus as an example to illustrate how age-specific incidence data can be used to quantify disease transmission and inform control interventions. Dengue is a relevant example because, despite being the most widely spread mosquito-transmitted virus, large gaps remain in our understanding of its global and local epidemiology(17). We present a model to estimate the transmission intensity of dengue from age-specific incidence data, and apply it to surveillance data from administrative units in four countries that suffer from endemic dengue transmission (Thailand, Colombia, Brazil and Mexico). We validate our estimates using serological data and show that they correlate well with known environmental drivers of dengue transmission at subnational level. Finally, we show how this approach could be used to guide the implementation of dengue control strategies such as vaccination.

## Results

We estimated the average forces of infection (FOI) over the last 20 years for 152 administrative level one units where age specific case data was available (Figure 1B): 72/76 provinces of Thailand, 28/32 departments of Colombia, 25/27 states of Brazil, and 27/31 states of Mexico. These administrative units comprise 90%, 99%, 96% and 91% of the population at risk of these countries respectively. We also estimated forces of infection for 211 municipalities (administrative level 2 units) of Colombia where at least 200 cases were reported over the period covered by the data.

**Figure 1:**
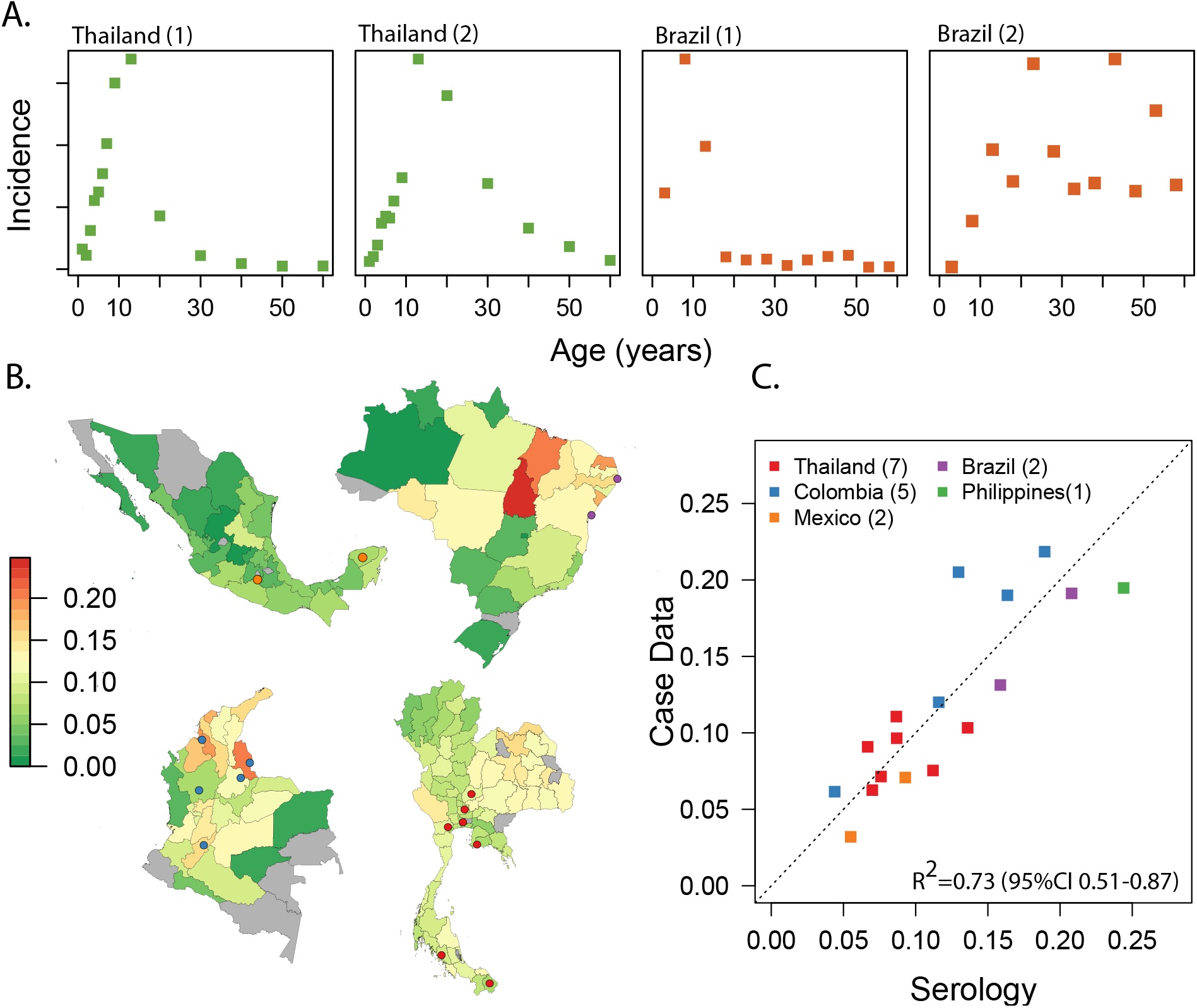
Estimating FOI from age-specific incidence data in Thailand, Colombia, Brazil and Mexico. Panel A shows examples of the age-specific incidence of dengue observed in two settings with very high endemic transmission (Thailand 1-Udon Thani, Thailand; Brazil 1 Pernambuco, Brazil) and two settings with lower and very low transmission (Thailand 2= Chiang Mai, Thailand; Brazil 2=Parana, Brazil). Panel B shows maps our estimates of the FOI for the four countries. Panel C correlates our estimates with those derived from age-stratified serological data (gold-standard) for 16 settings where we had both types of data (Thailand: Rayong (12); Bangkok (10); Ratchaburi (10); Lop Buri, Narathiwat, Trang, Ayuttayah(39). Philippines: Cebu (40). Brazil: Pernambuco (11); Salvador ((41)). Colombia (unpublished). Mexico: Morelos (42), Yucatan (43). The locations of the specific cities are shown in the maps in panel B.

The average force of infection (the rate at which susceptible individuals that will become infected in a year by any dengue serotype) was 0.096 in Thailand (95%CI 0.092-0.100), 0.132 (95%CI 0.128-0.136) in Colombia, 0.124 (95%CI 0.116-0.128) in Brazil, and 0.052 (95%CI 0.048-0.056) in Mexico. This implies that on average 9.6% of the susceptible population in Thailand gets infected every year by any of the circulating serotypes (1 – *e*^-*FOI*^). Similarly, 12% of the susceptible population of Colombia and Brazil, and 5% of the susceptible population of Mexico get infected yearly.

However, as expected, transmission intensity varied greatly within countries, ranging from 0.04 to 0.152 (coefficient of variation (CV) =0.27), between provinces of Thailand and between 0 and 0.204, 0.236 and 0.092 (CV= 0.37, 0.56 and 0.45), between departments/states of Colombia, Brazil and Mexico. Transmission was highest in the North East of Brazil (average FOI of 0.152) and in the Caribbean region of Colombia (average FOI of 0.156). There was also substantial heterogeneity within departments of Colombia. The mean CV for 15 departments where we had estimates for more than 5 municipalities was 0.4.

For the 17 locations where we had access to both age-stratified serological data (the gold standard) and case data, we found good correlation between the estimates of the force of infection derived from both data sources (R2=0.73, 95%CI 0.51-0.87, Figure 1C). In contrast, we found no correlation between recent incidence of dengue in these locations (the average yearly incidence over last 5 years where we had data) and the estimates of force of infection derived from serological data (R2=0.002, Figure S1).

Since estimates of transmission intensity derived from seroprevalence data are only available for a small number of locations, to further validate our method we also explored the association between our estimates of the force of infection for 211 Colombian municipalities (administrative level 2) with known environmental drivers of dengue transmission including temperature, elevation, a published metric of *Aedes aegypti* abundance (Figure 2) and population density (Table 2). Models were weighted by the number of cases used to estimate the FOI. On average, the force of infection increased by 0.006 (95% CI 0.004-0.007, R^2^=0.19) for each additional °C in temperature and, similarly, it decreased by 0.005 (95% CI 0.003, 0.006, R^2^=0.21) for each 100m increase in elevation. While population density was not associated with FOI estimates in unadjusted analyses, a 2-fold increase in density was associated with a 0.007 (95% 0.004-0.009) increase in FOI in the best fitting adjusted model. This model included elevation, population density and precipitation, and explained 35% (95%CI 23%-50%) of the variance.

**Figure 2:**
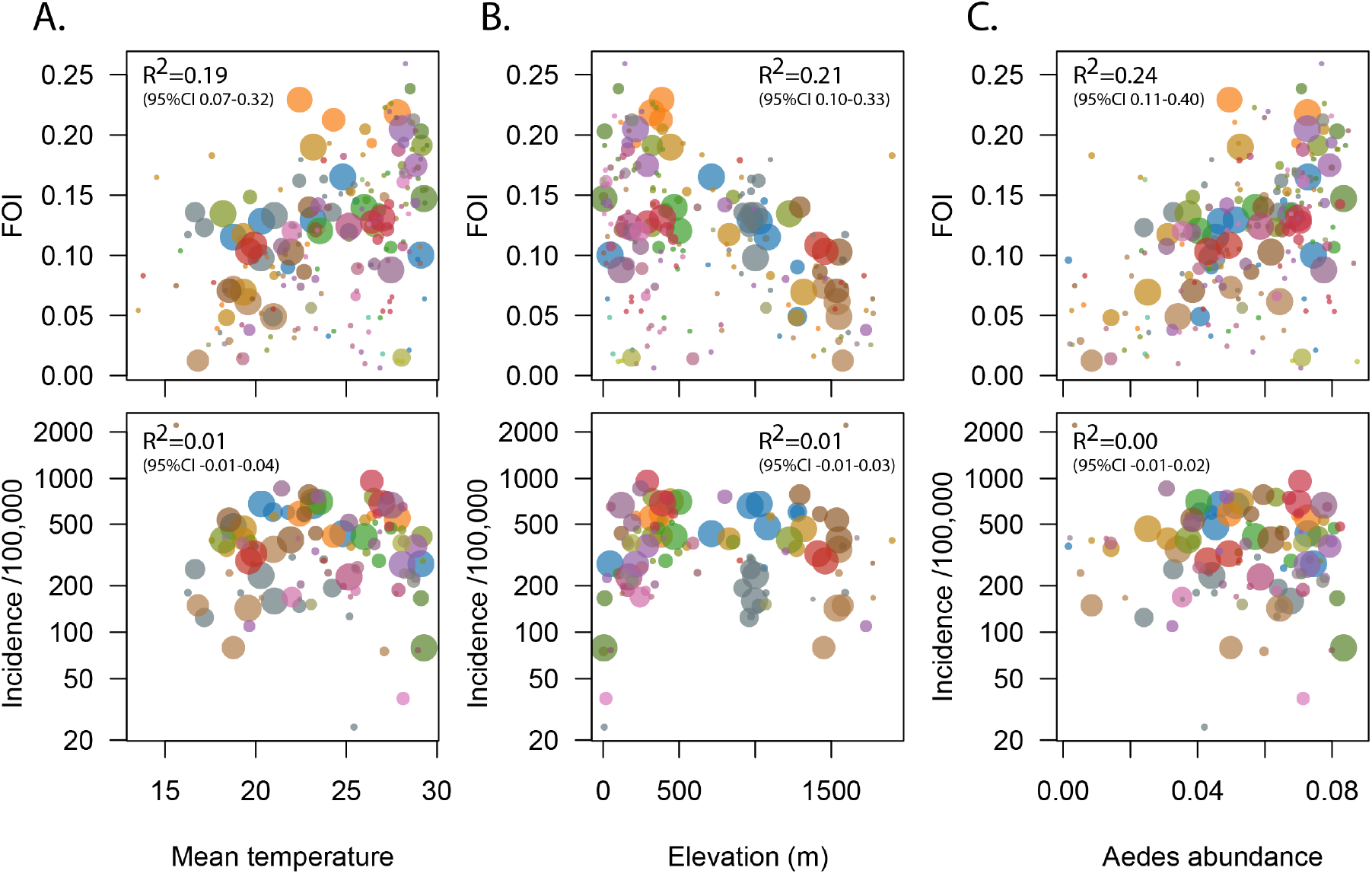
Correlation between estimates of the FOI and known environmental drivers of dengue transmission. Top panels show correlation between estimates of the FOI for 211 municipalities (administrative level 2) of Colombia and mean temperature **(A)**, elevation **(B)** and Aedes abundance **(C).** Size of symbols is proportional to the number of cases available to estimate the FOI. Bottom panels shows lack of correlation between environmental drivers and recent incidence, the most commonly used metric of transmission intensity. R^2^ values reported were obtained by fitting weighted linear regression models, with weights proportional to the number of cases used to derive the FOI estimate.

While we found strong associations between environmental variables and FOI, the recent incidence of dengue in these municipalities was not correlated with temperature, elevation or *Aedes aegypti* abundance (R2 0.01, 0.01, and 0.00 respectively, Table S1) We did find a negative association between population density and incidence, indicating a 6% (95%CI 2%-10%) decrease in log incidence for each 2-fold increase in population density.

## Application: Guiding dengue vaccination policy

The only available dengue vaccine (Dengvaxia®) has been licensed for use in children 9 years or older in 20 countries including Thailand, Brazil and Mexico. The WHO currently recommends confirmation (by virology or serology) of prior dengue infection at the individual level before vaccinating individuals, and therefore there is interest in identifying populations with high seropositivity to target pre-vaccination screening (18). In the absence of appropriate rapid serological assays that would allow implementing this individual screening strategy, an alternative that has been discussed is rolling out the vaccine in settings with 80% or greater seropositivity among 9-year olds. Using our estimates of the force of infection, we calculated the proportion of the population expected to be seropositive at age 9 years for each of the subnational units represented in our data. Our results show that only a small minority of locations in Colombia and Brazil have >80% seropositivity in this age group (Figure 3). The expected proportion seropositive at 9 years of age ranged between 0.35 and 0.75 in provinces of Thailand; 0.13 and 0.85 in departments of Colombia; 0 and 0.88 in states of Brazil; and 0.07 and 0.56 in states of Mexico. The seroprevalence was estimated to be high enough in only 2/28 Colombian departments, 4/25 Brazilian states and none of the 72 Thai provinces or 27 Mexican states. Furthermore, even within the 2 Colombian departments that met the 80% seroprevalence threshold, only 9/13 (70%) of the municipalities evaluated reached this level. This proportion would probably be much lower had we considered all the municipalities in these departments, as those excluded had, on average, lower FOI (Figure S2).

**Figure 3:**
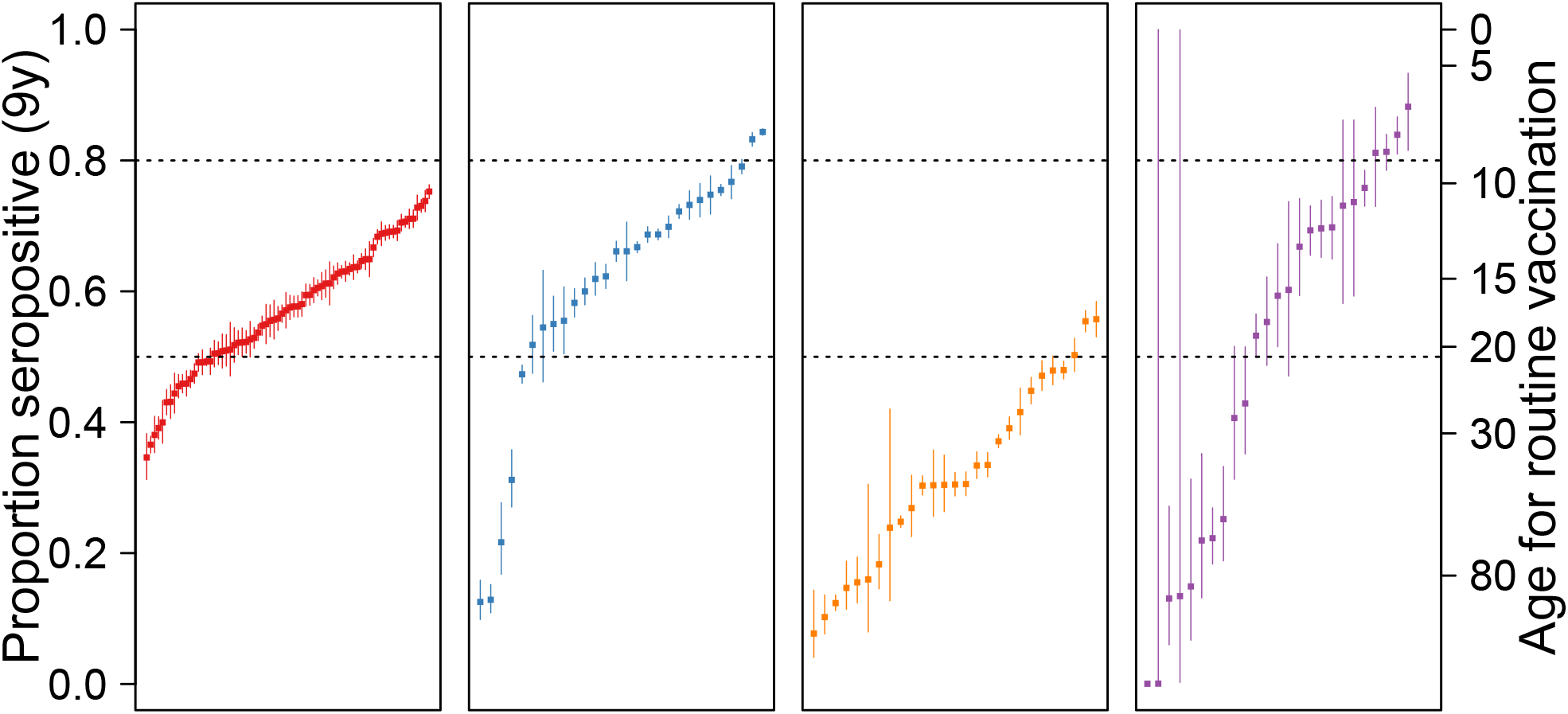
Guiding vaccination policy. Estimated dengue seroprevalence at 9 years of for administrative level 1 units of Thailand, Colombia, Brazil and Mexico. Expected seroprevalence of dengue among children aged 9 years, derived from the FOI estimates (see supplementary materials for more details), for administrative level 1 units of Thailand **(A),** Colombia **(B)**, Brazil (**C)** and Mexico **(D)**. Dashed lines indicate 50% and 80% seroprevalence levels. Therefore, units above the 80% line are those where, according to the WHO-SAGE recommendation from 2018, it might be reasonable target children aged 9 years old for vaccination. Units below the 50% line are those where vaccination of this age-group would not be recommended. The axis on the right of the plot indicates the minimum age-group that would need to be targeted in each location to ensure at least 80% seropositivity.

In locations where seroprevalence among 9 year olds was estimated to be less than 80%, we calculated the age-group that could be targeted to ensure a seroprevalence >80%. Our results suggest that, to comply with the WHO-SAGE recommendations, it would be necessary to target children 14 years of age or older in over 70% (108/152) of locations. Furthermore, in approximately 50% of the locations evaluated, the target vaccination need to be 18 years or older, precluding school-based vaccination strategies.

## Discussion

For most immunizing infections, age-stratified serological surveys are considered the gold-standard to measure the susceptibility of the population and quantify transmission. However, population-based serosurveys are expensive and labor intensive and therefore rarely available at a spatial or temporal resolutions useful to assess global and local epidemiologic patterns or to target control interventions. We developed a model to estimate the force of infection of dengue from age-specific incidence data, routinely collected by most surveillance systems, and generated country-wide subnational estimates for Thailand, Colombia, Brazil and Mexico.

Our findings are consistent with large variation in dengue transmission between and within countries. Spatial heterogeneity was substantial not only between, but within administrative level 1 units, indicating that decisions of where to deploy control interventions, including vaccines, should be made at least at the municipal/district (administrative level 2) level. Transmission intensity was highest in northeastern Brazil, northern Colombia and eastern Thailand. Not surprisingly, northeastern Brazil was also the region that reported the highest incidence of births with microcephaly during the 2015/2016 epidemic of Zika virus, transmitted by the same mosquito vector (*Aerdes aegypti)*(19). While this heterogeneity is probably driven by multiple environmental, socio-economic and demographic factors, our results suggest that as much as 35% of the variance may be explained by differences in temperature/elevation and population density alone.

Most surveillance systems use case counts or incidences to describe temporal and spatial trends of communicable diseases. Our results underscore the extent to which, for immunizing diseases in endemic circulation, recent incidence may be poor metric of transmission and may be misleading when ranking spatial units (Figure S3). Immunity of the population in high transmission settings reduces the number of individuals that are susceptible to infection. As a result, incidence in places where transmission intensity is lower, but people remain susceptible for a longer period of time, may be roughly equivalent to that in higher transmission intensity areas. Metrics such as the force of infection, that quantify the risk among the susceptible population, better reflect the underlying transmission potential. Since FOI estimates are derived from the age distribution of incidence, and not from the aggregate counts, they are more robust to differences in surveillance efficiency and can be obtained with relatively small numbers of yearly cases. These findings raise concerns about the vaccination policies that were decided in some settings based on recent incidence, in the absence of seroprevalence data. In particular, the decision to deploy the vaccine in the Brazilian state of Paraná is highly questionable, as estimates suggest that seroprevalence is likely to be well below the recommended threshold for vaccination, even among adults (20).

There are several limitations of using age-specific surveillance data to estimate transmission parameters of dengue. Of concern are age specific differences in the probability of clinical disease upon infection or in health care utilization that could bias estimates derived from case data either up or down depending on the specific bias. Differences in reporting practices between and within countries could also limit our capacity to reconstruct transmission history from case-data. Our model also makes several assumptions that may be questionable. In particular, it assumes that the age distribution of cases represents the age-distribution of secondary infections, thus ignoring the potential contribution of primary, tertiary and quaternary infections. It also assumes that risk is not age dependent, even though there is some evidence that suggests that certain age-groups may be at increased risk(21-23). Finally, it assumes equal circulation of all serotypes, despite the known dominance of specific serotypes for extended periods of time in the Americas.(21,24) Despite these simplifications, validation of our estimates using age-stratified serological data from 17 locations is very encouraging, as is the good correlation with known drivers of dengue transmission. While further validation is desirable, it is important to note that some discrepancy between our estimates and those derived from seroprevalence data is expected as the two sources of data do not represent exactly the same period of time and location. For example, most of the serosurveys available were conducted in specific urban centers, while case-data represents the full administrative unit.

Targeting control interventions against dengue and other communicable diseases requires good understanding of when and where transmission is occurring. Careful analyses of age-specific incidence data, collected by surveillance systems at high temporal and spatial resolution, can provide very useful information to characterize transmission and target control interventions at spatial scales at which serological data is rarely available. While here we present average forces of infection, these methods can also be used to reconstruct changes in transmission over time (25,26). Open-access to age-specific incidence data would greatly enrich and enhance existing efforts to quantify trends in the global burden of disease.

## Methods

### Data used

We used data on the yearly age-specific number of dengue and dengue hemorrhagic fever cases for administrative level 1 units of Thailand, Colombia, Brazil and Mexico (27-30) as well as administrative level 2 units from Colombia. We also used population data from each administrative unit analyzed, available from the national statistical office of each country. Information on the type, source and years of data used are provided in Table 1.

**Table 1:**
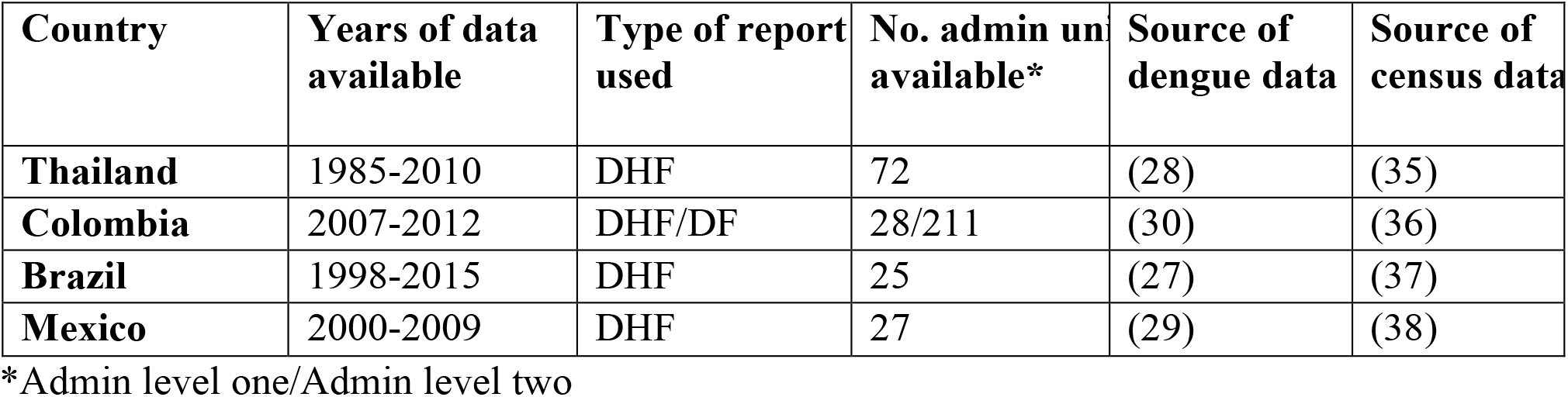
Data sources used

**Table 2:**
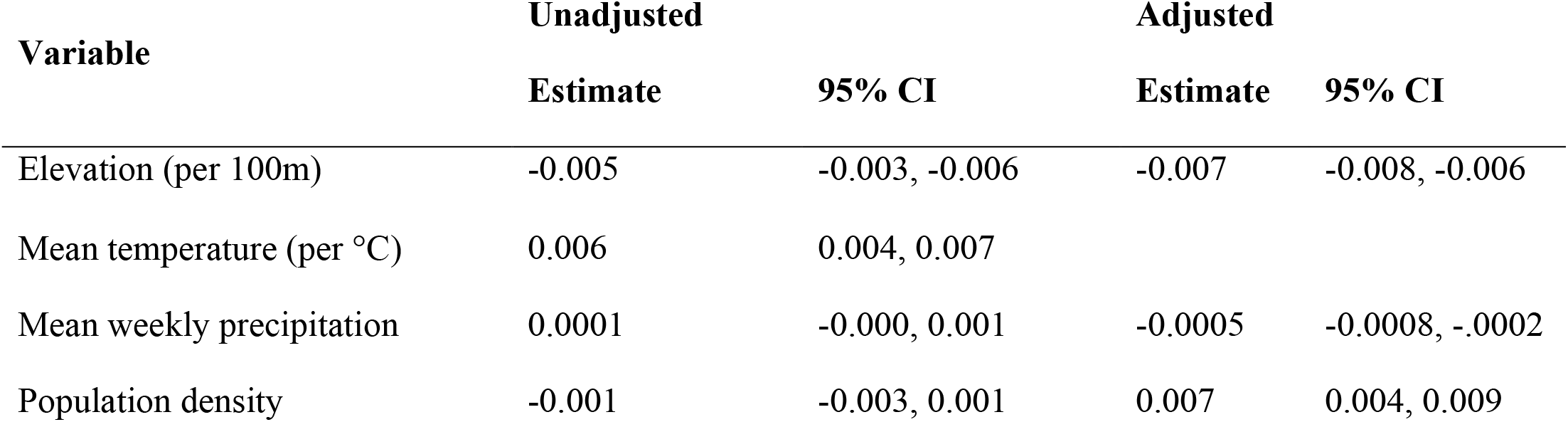
Association between environmental variables and dengue force of infection for 211 municipalities in Colombia

### Statistical analyses

We estimated the average force of infection of dengue, over the last 20 years, for each administrative unit for which we had available data. The force of infection (λ) is a metric used to characterize the transmission intensity in a specific setting and estimates the per capita rate at which susceptible individuals are infected. Methods to estimate transmission intensity from age-specific incidence data have been previously used to reconstruct the transmission history of measles and dengue(7,26). Briefly, these methods rely on the fact that, for immunizing infections, accumulation of immunity shapes the age distribution of future cases. In settings with high endemic transmission, incident cases are expected to be concentrated in younger age groups, as adults are likely to be already immune (Figure 1A). In contrast, in places where there is less population immunity, the age distribution of cases is more likely to resemble the age distribution of the population itself, with cases in in both children and adult populations.

Methods to estimate dengue forces of infection from case data have been applied to settings where dengue is thought to be close to endemic circulation (26,31). These methods generally rely on the cumulative incidence proportion, and therefore assume that all individuals are infected by dengue at some point in their lifetime. They also often assume that the distribution of cases (of dengue hemorrhagic fever cases in particular) is representative of the distribution of secondary cases. Here, we extend these methods to accommodate settings where transmission hazards are lower or where dengue may have been more recently introduced. We do this by modeling directly the age-specific incidence of cases, rather than the cumulative incidence proportion. Details of our model are provided in the supplementary material and code to implement the model is available at https://github.com/isabelrodbar/dengue_foi.

We fit all models in a Bayesian Markov chain Monte Carlo (MCMC) framework using the RStan package in R, using wide priors (Normal distribution with mean 0 and standard deviation of 1000). We simulated four independent chains, each of 30000 iterations and discarded the first 10000 iterations as warm-up. We assessed convergence visually and using Rubin’s R statistic. We obtained 95% credible intervals from the 2.5% and 97.5% percentiles of the posterior distributions.

### Validation and sensitivity analyses

We validated our estimates of the force of infection by comparing them to those obtained from age-stratified serological data (the gold-standard) for 17 locations where we had both serologic and age-specific case data.

Since dengue transmission is known to be highly spatially heterogeneous, we also correlated our administrative level 2 estimates for Colombia with known environmental drivers of dengue transmission: temperature, elevation, population density and a published composite metric of *Aedes aegypti* abundance(32).

As stated above, a key assumption of this model is that the age distribution of cases represents the age distribution of secondary infections. Since data from Thailand has consistently suggested that the majority of dengue hemorrhagic fever (DHF) cases arise from secondary infections(33), we limited our analysis to reports of DHF where possible (Thailand, Brazil and Mexico). However, for Colombia we used combined DF/Severe dengue data because the severe dengue data alone was too sparse. To assess the impact of the data type, we performed sensitivity analyses including all dengue cases, as it is known that many surveillance systems do not differentiate between types of dengue disease.

### Application: Guiding dengue vaccination policy

The first dengue vaccine has been licensed for use in children over 9 years of age in 20 countries. Due to uncertainty regarding the vaccine’s benefits and risks in individuals who haven’t been previously infected by dengue, the WHO’s scientific advisory group of experts (SAGE) committee recommended in April 2016 that this vaccine only be used in settings with known high endemicity, defined as places where seroprevalence is greater than 70% in the target vaccination age-group (34), and should not be used in places where seroprevalence is under 50%. This recommendation was later revised, and the WHO now recommends that individuals should be tested for dengue antibodies prior to vaccination, and the vaccine should only be given to individuals who have been infected by dengue in the past(18). In the absence of appropriate serological assays that would allow for pre-vaccination screening, an alternative that has been discussed is deploying the vaccine in settings were seroprevalence is 80% or greater. These recommendations pose challenges to countries wanting to implement the vaccine, as they require detailed knowledge of the epidemiology of dengue. Specifically, they require knowledge of the population seroprevalence against dengue at subnational levels, even though such data is not available.

In order to provide information useful to countries considering deploying the vaccine according to the WHO recommendations, we used our estimates of the force of infection to calculate the proportion of the population expected to be seropositive at age 9 years for of the subnational units represented in our data (See supplementary material). We also estimated the minimum age group expected to have a seroprevalence of 80% or greater. In collaboration with the MRC center for Outbreak Analysis and Modelling, these estimates were made available online in June 2017 (https://mrcdata.dide.ic.ac.uk/_dengue/dengue.php) as a tool to help countries deciding where to target vaccination.

## Acknowledgements

We thank the Ministry of Public Health (Thailand), the Instituto Nacional de Salud (INS, Colombia), the Ministerio da Saude (Brazil) and the Direccion General de Epidemiología. Sistema Nacional de Vigilancia Epidemiológica (Sinave, Mexico), for making the data necessary for these analyses publicly available. We also thank Sompong Vongpunsawad for sharing the raw data of the serological studies conducted in Thailand. We thank Neil Ferguson, Natsuko Imai and Wes Hinsley for valuable input and for making these estimates publicly available through the Global Dengue Transmission Map website.

## Supplementary material captions

**Text S1**: Supplementary methods and results

## Methods

### Model

#### i. Estimating the force of infection

The force of infection (λ) is a measure used to characterize infection hazard in a given setting and estimates the per capita rate of acquisition of infection by susceptible individuals. Methods to estimate the force of infection from age-stratified serosurveyshave been described extensively elsewhere^1^. When serosurvey data is not available, age-stratified incidence data can also be used to estimate forces of infection, as age patterns of disease depend on the age distribution of susceptible individuals in the population ^2^.

Methods have been described to estimate dengue forces of infection using case data in settings where dengue has been in endemic circulation^3^. These methods rely on the cumulative incidence proportion, and therefore assume that all individuals are infected by degue at some point in their lifetimes. They also assume that the age distribution of incident cases is representative of the age distribution of secondary infections, the incidence.

Here, we adapt these methods to accommodate settings where transmission hazards are lower or where dengue may have been more recently introduced. We do this by modeling directly the age-specific incidence of cases, rather than the cumulative incidence proportion.

The fraction of the population susceptible to all dengue serotypes at age a and t, x(a,t) is given by

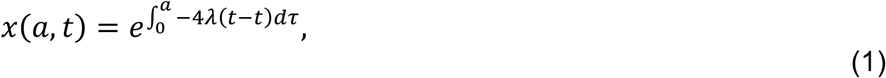

where λ(t) is average force of infection per serotype at time *t* and 4λ(t) is the total force of infection assuming four circulating serotypes. The proportion of individuals of age a who have been infected with only serotype at time *t*, but are still susceptible to all other serotypes is denoted z_1_(a,t) and is given by:

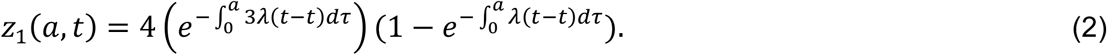

Assuming that the age-specific incidence of cases is representative of the distribution of secondary infections, the expected incidence rate among individuals age *a* at time *t* is given by

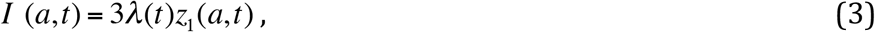

and the expected reported number of cases is

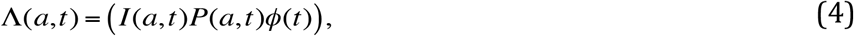

where P(a,t) is the size of the population aged a at time t, and Φ(t) represents a time specific reporting rate.

#### ii. Likelihood and estimation

Assuming that the observed age specific case counts C(a,t) follow a Poisson distribution, the likelihood of the data can be expressed as

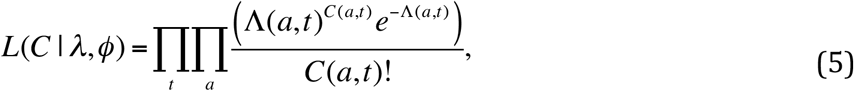

We fit the model in a Bayesian Markov chain Monte Carlo (MCMC) framework using the RStan package in R ^4 5^. Both the annual hazards of infection (λ) and the reporting rates (Φ) were estimated on a logit scale using wide priors (Normal distribution with mean 0 and standard deviation of 1000). We simulated four independent chains, each of 30000 iterations and discarded the firs 10000 iterations as warm-up. We assessed convergence visually and using Rubin’s R statistic. We obtained 95% credible intervals from the 2.5% and 97.5% percentiles of the posterior distributions.

A limitation of this approach is that, due to the large number (often in the thousands) that are characteristic of the data for some settings, the estimated confidence intervals produced are extremely narrow and do not reflect the underlying uncertainty adequately. The observed counts can also be assumed to follow a negative binomial distribution to account for some overdispersion.

#### iii. Parameters estimated

Since it is known that the force of infection has varied substantially over time in many of the settings considered, we allowed λ(t) to vary as a function of time. To limit the number of parameters estimated, we assumed constant λ(t) for periods of 20 years. Thus, if for a given setting we were estimating hazards for the period 1935-2015, we assumed piecewise-constant λ(t)s for the periods 1935-1954, 1955-1974, 1975-1994, 1995-2015. Given the objective of this study was to characterize recent transmission in endemic settings, we focused our results on the estimate of the average λ(t) for the most recent 20-year period. In the main text we focus on reporting the total force of infection (4λ(t)).

To account for the large variation in yearly dengue incidence, our model also included yearly “reporting rates” Φ(t). These reporting rates not only capture variations in reporting itself, but also variation in the symptomatic:asymptomatic ratio of dengue infections.

#### iv. Estimating the proportion seropositive at a given age

Using our estimates of the average force of infection, we estimated the proportion of individuals expected to be seropositive by age y(x) as:

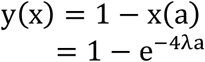

Where x(a) is the proportion of the population susceptible at age a and λ is the average force of infection per serotype (assuming four serotypes circulating). Since the vaccine has been registered for use in children 9 years of age or older, we report the proportion of individuals expected to be seropositive by age 9 years for each of the settings.

#### v. Estimating the minimum age to achieve a given level of seropositivity

Given that the WHO initially recommended this vaccine in places where at least 70% of the target age-group is seropositive, we estimated the minimum age at which this level of seropositivity is expected for each of the settings.

For a given level of transmission λ it is possible to estimate the minimum age (A) at which a given level of seropositivity (s) is expected as:

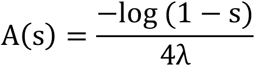

## Results

### Supplementary results

**Figure S1:**
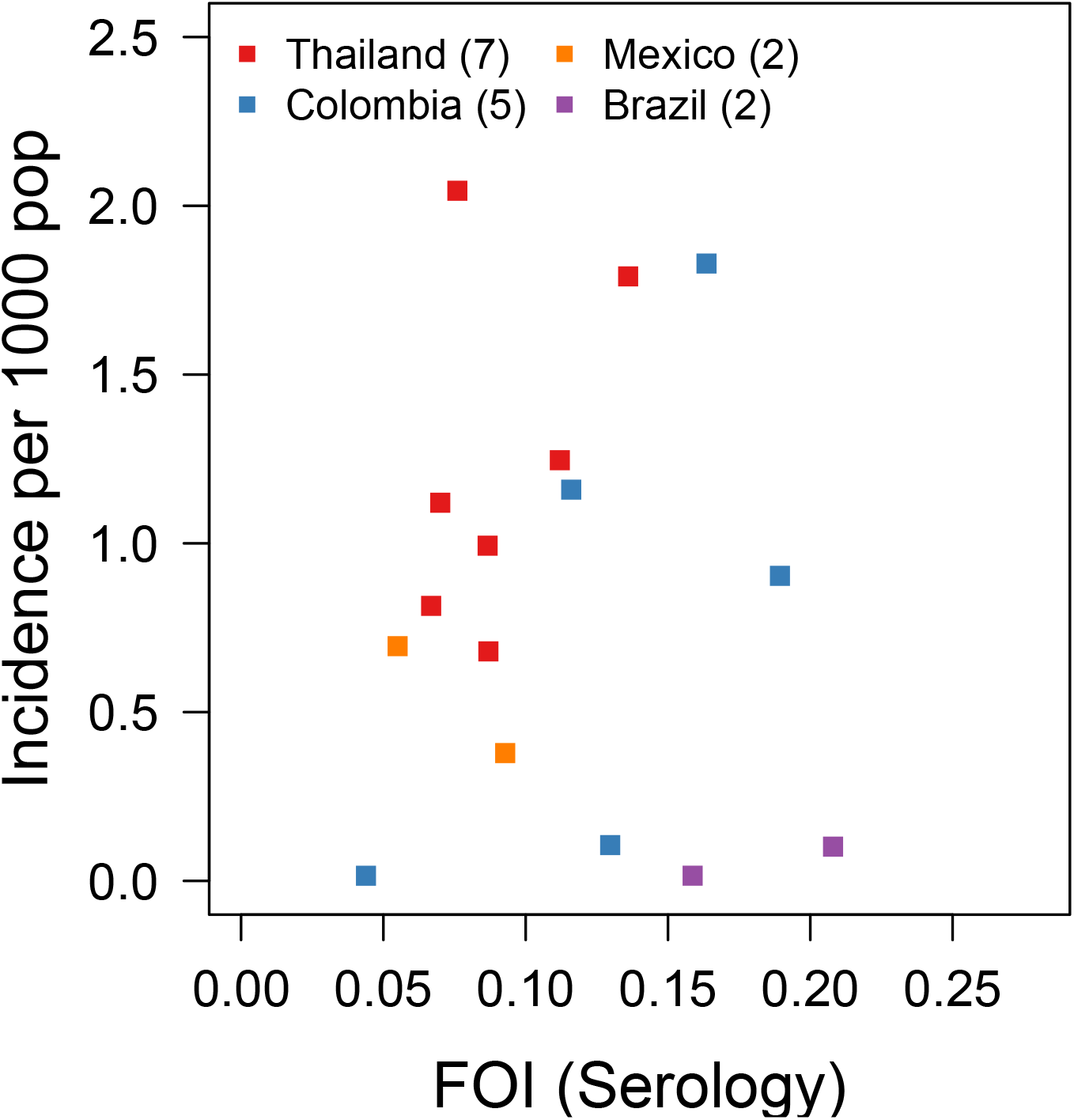
Correlation between the FOI estimates obtained from serological data and the recent incidence of DHF in 16 locations where we had both types of data (R^2^=0.002).

**Figure S2:**
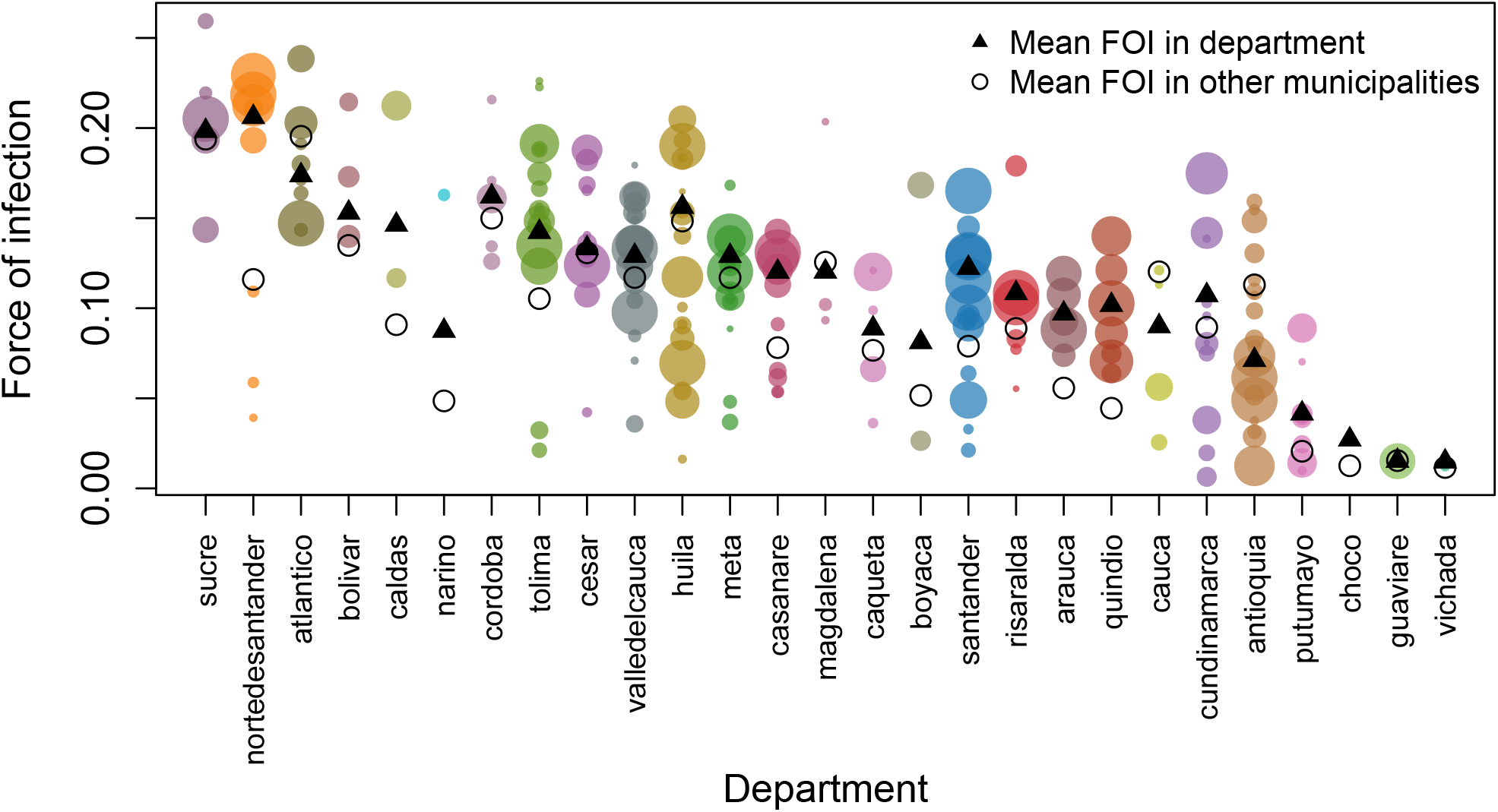
Heterogeneity in FOI within administrative level 1 units of Colombia. Filled circles show the estimates for each municipality (administrative level 2 unit) within the department that reported >200 cases. Hollow circles indicate the mean force of infection for those municipalities that reported <200 cases. Size of circles is proportional to the number of cases available to estimate the FOI. Triangles indicate the mean estimate for each department.

**Figure S3:**
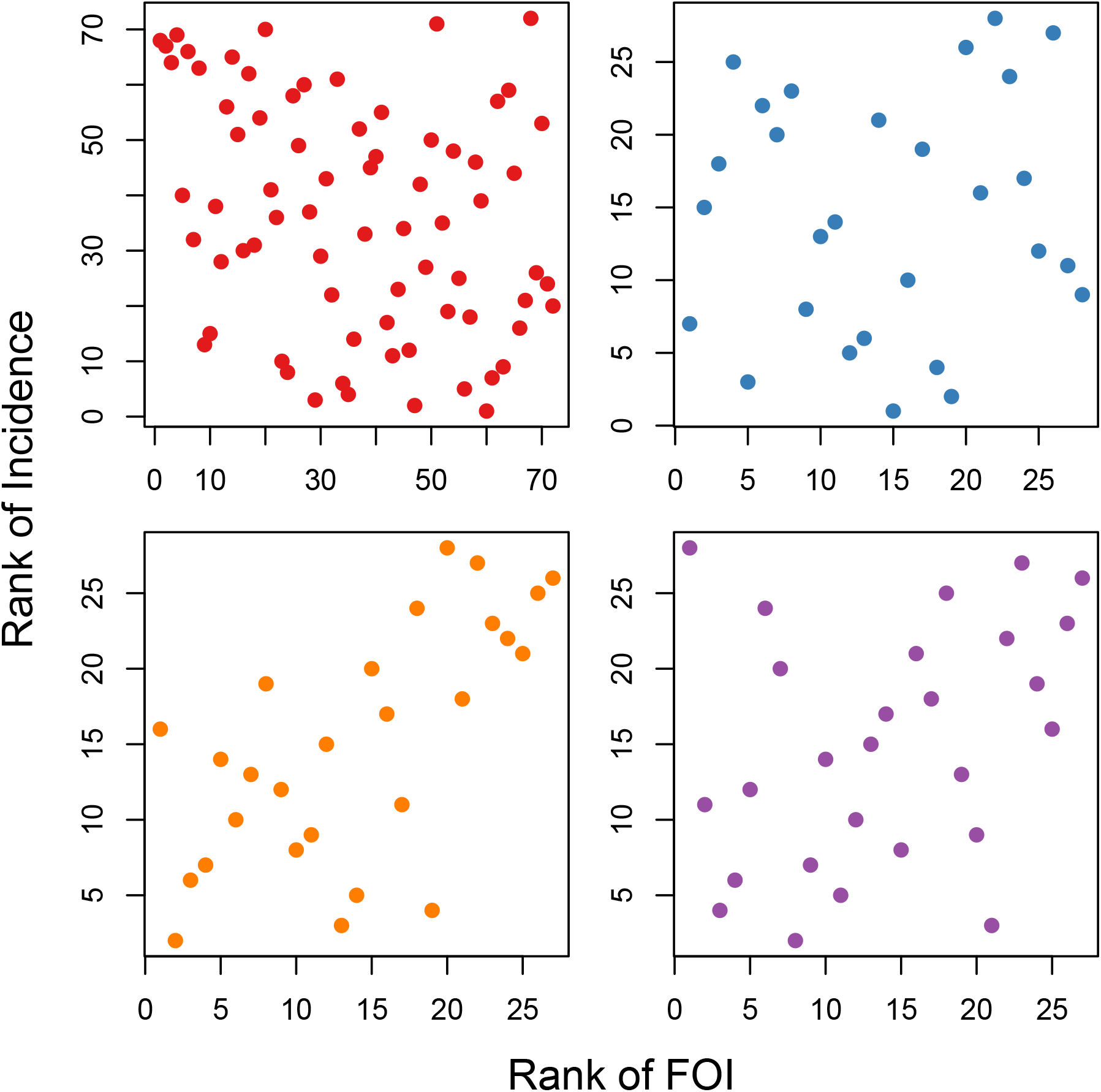
Ranking of administrative level-1 units of Thailand, Colombia, Mexico and Brazil based on FOI estimates (x-axis) and incidence in the past 5-years (y-axis).

**Table S1:** Association between environmental variables and the log incidence of dengue (over the last 5 years) for 211 municipalities in Colombia.

| **Variable** | **Unadjusted** |  | **Adjusted\*** |  |
| --- | --- | --- | --- | --- |
|  | **Estimate** | **95% CI** | **Estimate** | **95% CI** |
| Elevation (per 100m) | -0.02 | -0.03, 0 | 0.01 | -0.01, 0.03 |
| Mean temperature (per C) | 0.01 | 0 , 0.05 |  |  |
| Mean weekly precipitation | 0 | 0, 0 | -0.002 | -0.01, 0 |
| Population density | -0.04 | -0.07, -0.01 | -0.06 | -0.10, -.02 |

\* R2 of adjusted model: 0.04

### Estimates

**Table S2:**
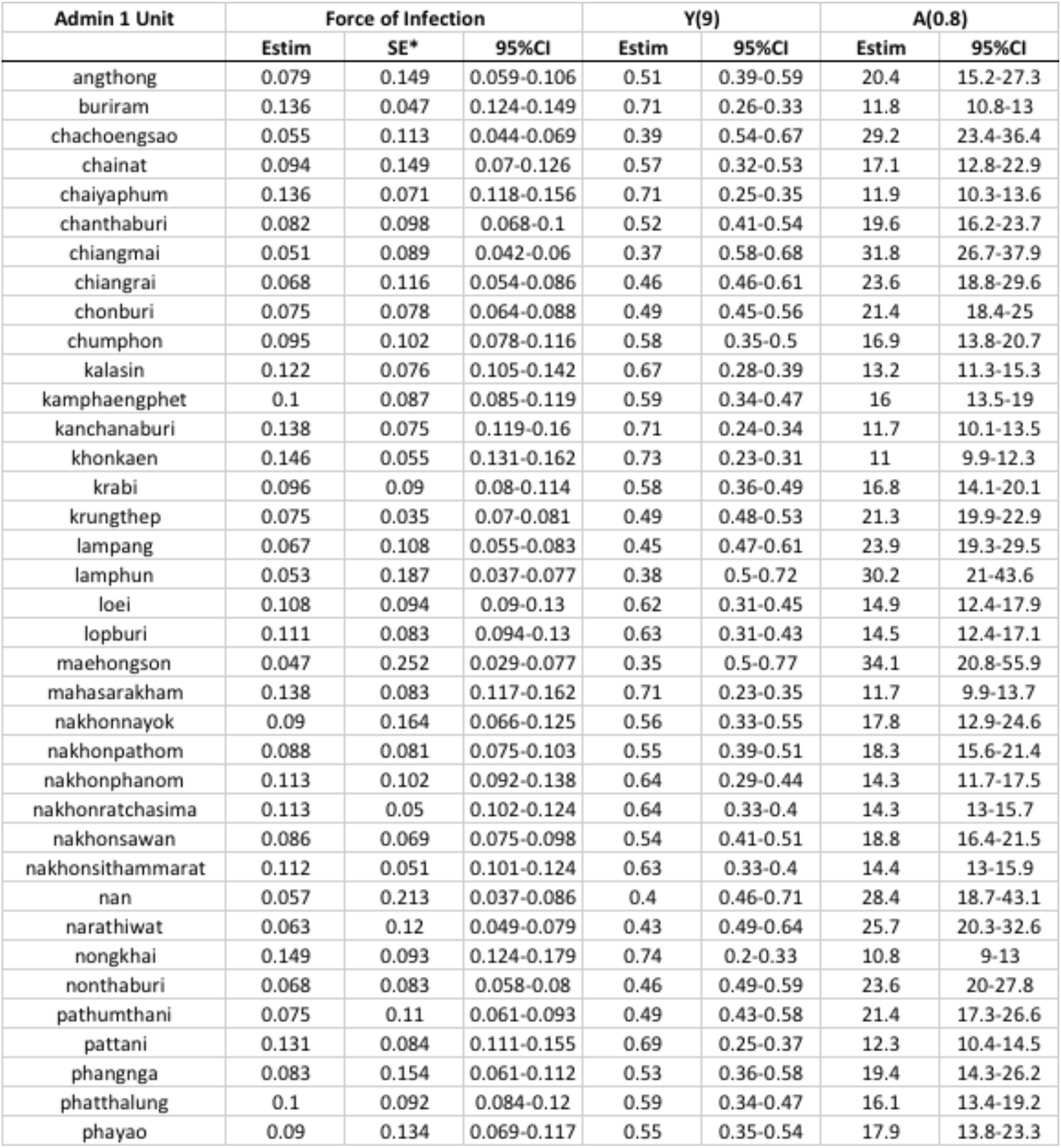
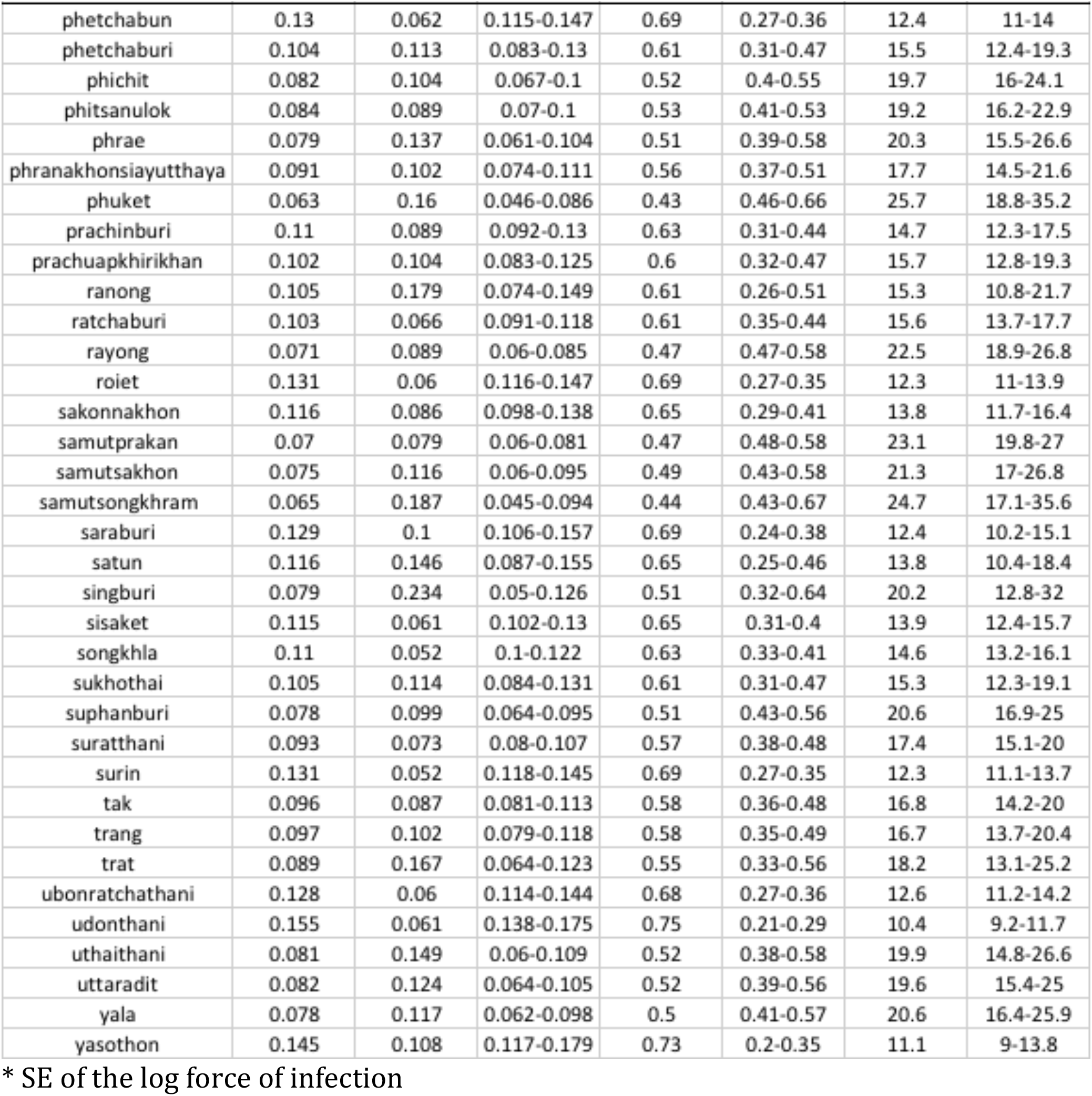
Estimates for administrative level 1 units of Thailand.

**Table S3:** Estimates for administrative level 1 units of Brazil.

| Admin 1 Unit | Force of Infection |  |  | Y(9) |  | A(0.8) |  |
| --- | --- | --- | --- | --- | --- | --- | --- |
|  | Estim | SE* | 95%CI | Estim | 95%CI | Estim | 95%CI |
| alagoas | 0.102 | 0.757 | 0.023-0.452 | 0.6 | 0.19-0.98 | 15.7 | 3.6-69.3 |
| amapa | 0.016 | 8.179 |  | 0.13 | 0-1 | 100.5 |  |
| amazonas | 0 | 61.082 |  |  |  |  |  |
| bahia | 0.131 | 0.212 | 0.087-0.199 | 0.69 | 0.54-0.83 | 12.3 | 8.1-18.6 |
| ceara | 0.132 | 0.247 | 0.082-0.215 | 0.7 | 0.52-0.86 | 12.2 | 7.5-19.7 |
| espiritosanto | 0.058 | 0.673 | 0.016-0.217 | 0.41 | 0.13-0.86 | 27.8 | 7.4-103.8 |
| goias | 0.028 | 0.445 | 0.012-0.067 | 0.22 | 0.1-0.45 | 57.5 | 24-137.5 |
| maranhao | 0.203 | 0.198 | 0.138-0.299 | 0.84 | 0.71-0.93 | 7.9 | 5.4-11.7 |
| matogrosso | 0.133 | 0.27 | 0.078-0.226 | 0.7 | 0.51-0.87 | 12.1 | 7.1-20.6 |
| matogrossosul | 0.032 | 0.675 | 0.009-0.121 | 0.25 | 0.07-0.66 | 49.9 | 13.3-187.5 |
| minasgerais | 0.09 | 0.384 | 0.042-0.19 | 0.55 | 0.32-0.82 | 18 | 8.5-38.2 |
| para | 0.1 | 0.442 | 0.042-0.238 | 0.59 | 0.32-0.88 | 16.1 | 6.8-38.3 |
| paraiba | 0.123 | 0.416 | 0.054-0.277 | 0.67 | 0.39-0.92 | 13.1 | 5.8-29.7 |
| parana | 0.028 | 1.146 | 0.003-0.26 | 0.22 | 0.03-0.9 | 58.5 | 6.2-552.8 |
| pernambuco | 0.158 | 0.163 | 0.115-0.217 | 0.76 | 0.64-0.86 | 10.2 | 7.4-14.1 |
| piaui | 0.146 | 0.837 | 0.028-0.752 | 0.73 | 0.22-1 | 11 | 2.1-56.9 |
| riodejaneiro | 0.084 | 0.189 | 0.058-0.122 | 0.53 | 0.41-0.67 | 19.1 | 13.2-27.6 |
| riograndedonorte | 0.186 | 0.179 | 0.131-0.265 | 0.81 | 0.69-0.91 | 8.6 | 6.1-12.3 |
| riograndedosul | 0.018 | 1.723 | 0.001-0.525 | 0.15 | 0.01-0.99 | 89.7 | 3.1-2629.6 |
| rondonia | 0.148 | 0.805 | 0.031-0.718 | 0.74 | 0.24-1 | 10.9 | 2.2-52.7 |
| roraima | 0.016 | 1.667 | 0.001-0.408 | 0.13 | 0.01-0.97 | 103.4 | 3.9-2716 |
| saopaulo | 0.062 | 0.524 | 0.022-0.174 | 0.43 | 0.18-0.79 | 25.9 | 9.3-72.3 |
| sergipe | 0.185 | 0.498 | 0.07-0.492 | 0.81 | 0.47-0.99 | 8.7 | 3.3-23 |
| tocantins | 0.238 | 0.48 | 0.093-0.609 | 0.88 | 0.57-1 | 6.8 | 2.6-17.4 |

\* SE of the log force of infection

**Table S4:**
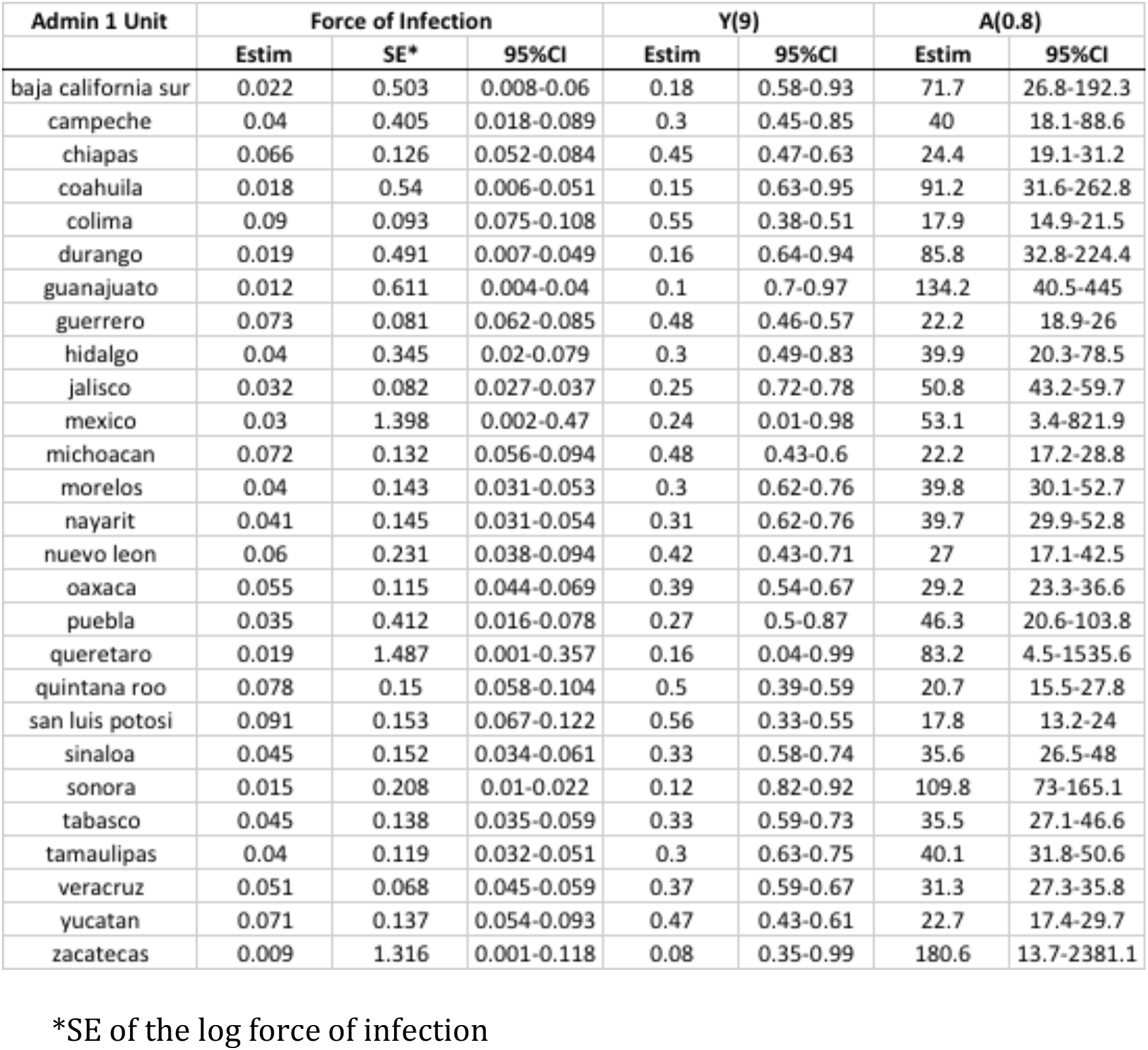
Estimates for administrative level 1 units of Mexico.

**Table S5:** Estimates for administrative level 1 units of Colombia.

| Admin 1 Unit | Force of Infection |  |  | Y(9) |  | A(0.8) |  |
| --- | --- | --- | --- | --- | --- | --- | --- |
|  | Estim | SE* | 95%CI | Estim | 95%CI | Estim | 95%CI |
| antioquia | 0.071 | 0.082 | 0.061-0.084 | 0.47 | 0.47-0.58 | 22.6 | 19.2-26.5 |
| arauca | 0.097 | 0.119 | 0.077-0.122 | 0.58 | 0.33-0.5 | 16.6 | 13.1-20.9 |
| atlantico | 0.174 | 0.067 | 0.152-0.198 | 0.79 | 0.17-0.25 | 9.3 | 8.1-10.6 |
| bolivar | 0.153 | 0.168 | 0.11-0.213 | 0.75 | 0.15-0.37 | 10.5 | 7.6-14.6 |
| boyaca | 0.081 | 0.251 | 0.05-0.133 | 0.52 | 0.3-0.64 | 19.8 | 12.1-32.5 |
| caldas | 0.146 | 0.123 | 0.115-0.186 | 0.73 | 0.19-0.36 | 11 | 8.6-14 |
| caqueta | 0.089 | 0.234 | 0.056-0.14 | 0.55 | 0.28-0.6 | 18.1 | 11.5-28.7 |
| casanare | 0.12 | 0.084 | 0.102-0.142 | 0.66 | 0.28-0.4 | 13.4 | 11.4-15.8 |
| cauca | 0.09 | 0.284 | 0.052-0.157 | 0.56 | 0.24-0.63 | 17.9 | 10.3-31.2 |
| cesar | 0.133 | 0.092 | 0.111-0.16 | 0.7 | 0.24-0.37 | 12.1 | 10.1-14.5 |
| choco | 0.027 | 0.569 | 0.009-0.083 | 0.22 | 0.47-0.92 | 59.3 | 19.4-181 |
| cordoba | 0.162 | 0.15 | 0.121-0.217 | 0.77 | 0.14-0.34 | 9.9 | 7.4-13.3 |
| cundinamarca | 0.107 | 0.136 | 0.082-0.14 | 0.62 | 0.28-0.48 | 15 | 11.5-19.6 |
| guaviare | 0.015 | 0.36 | 0.008-0.031 | 0.13 | 0.76-0.93 | 105.2 | 52-212.8 |
| huila | 0.156 | 0.05 | 0.142-0.172 | 0.75 | 0.21-0.28 | 10.3 | 9.3-11.4 |
| laguajira | 0.15 | 0.146 | 0.112-0.199 | 0.74 | 0.17-0.36 | 10.8 | 8.1-14.3 |
| magdalena | 0.12 | 0.245 | 0.074-0.194 | 0.66 | 0.17-0.51 | 13.4 | 8.3-21.6 |
| meta | 0.129 | 0.065 | 0.114-0.147 | 0.69 | 0.27-0.36 | 12.5 | 11-14.2 |
| narino | 0.087 | 0.479 | 0.034-0.224 | 0.54 | 0.13-0.74 | 18.4 | 7.2-47.1 |
| nortedesantander | 0.206 | 0.034 | 0.193-0.22 | 0.84 | 0.14-0.18 | 7.8 | 7.3-8.3 |
| putumayo | 0.042 | 0.34 | 0.021-0.081 | 0.31 | 0.48-0.83 | 38.7 | 19.9-75.4 |
| quindio | 0.102 | 0.112 | 0.082-0.127 | 0.6 | 0.32-0.48 | 15.8 | 12.7-19.7 |
| risaralda | 0.108 | 0.1 | 0.089-0.132 | 0.62 | 0.31-0.45 | 14.9 | 12.2-18.1 |
| santander | 0.122 | 0.046 | 0.112-0.134 | 0.67 | 0.3-0.37 | 13.2 | 12-14.4 |
| sucre | 0.198 | 0.069 | 0.173-0.227 | 0.83 | 0.13-0.21 | 8.1 | 7.1-9.3 |
| tolima | 0.142 | 0.059 | 0.127-0.16 | 0.72 | 0.24-0.32 | 11.3 | 10.1-12.7 |
| valledelcauca | 0.129 | 0.045 | 0.118-0.141 | 0.69 | 0.28-0.35 | 12.5 | 11.4-13.6 |
| vichada | 0.015 | 0.507 | 0.006-0.04 | 0.13 | 0.7-0.95 | 108.1 | 40-292.1 |

\*SE of the log force of infection

### Sensitivity analyses

In general, using data from all cases produces estimates that are lower than those obtained when using data from more severe cases (DHF). This is likely due to the fact that milder forms of the disease probably arise from a mixture of primary – quaternary infections, and not just secondary infections. Thus, inference made for places where all-cases are reported is likely to be conservative.

**Figure S4 figure supplement 2:**
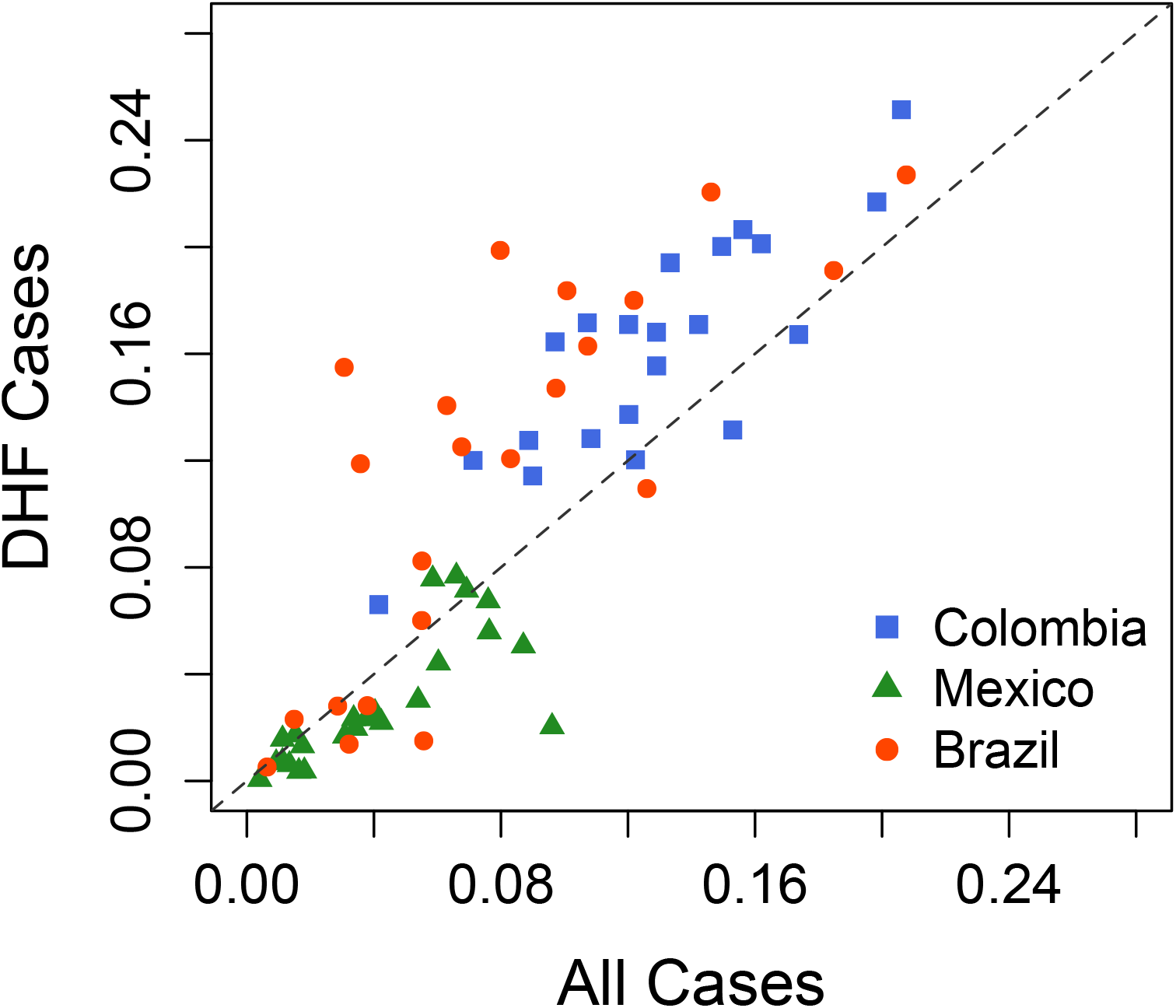
Comparison of estimates obtained when using all data vs. data from severe infections only.

### Selected model fits

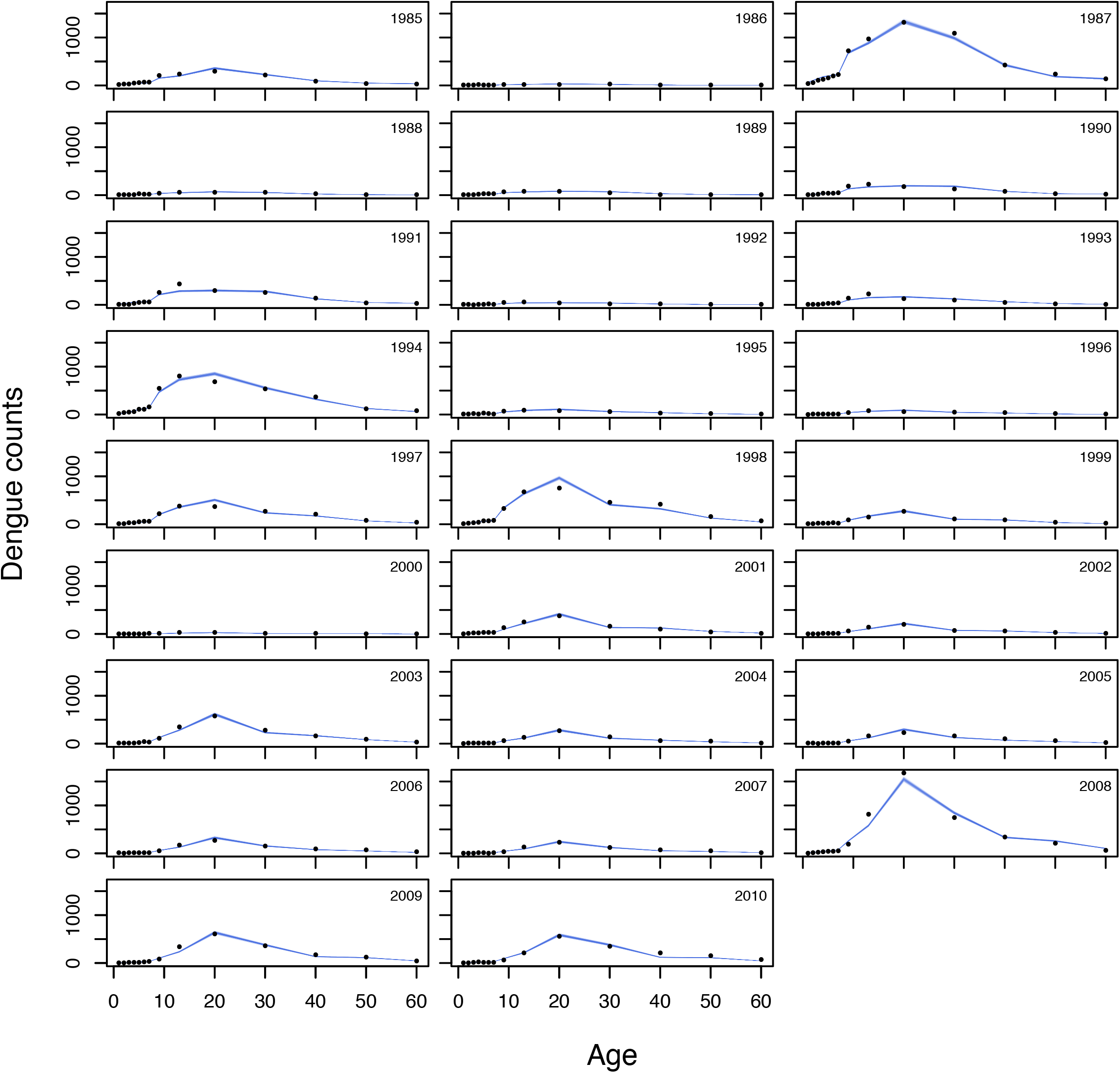

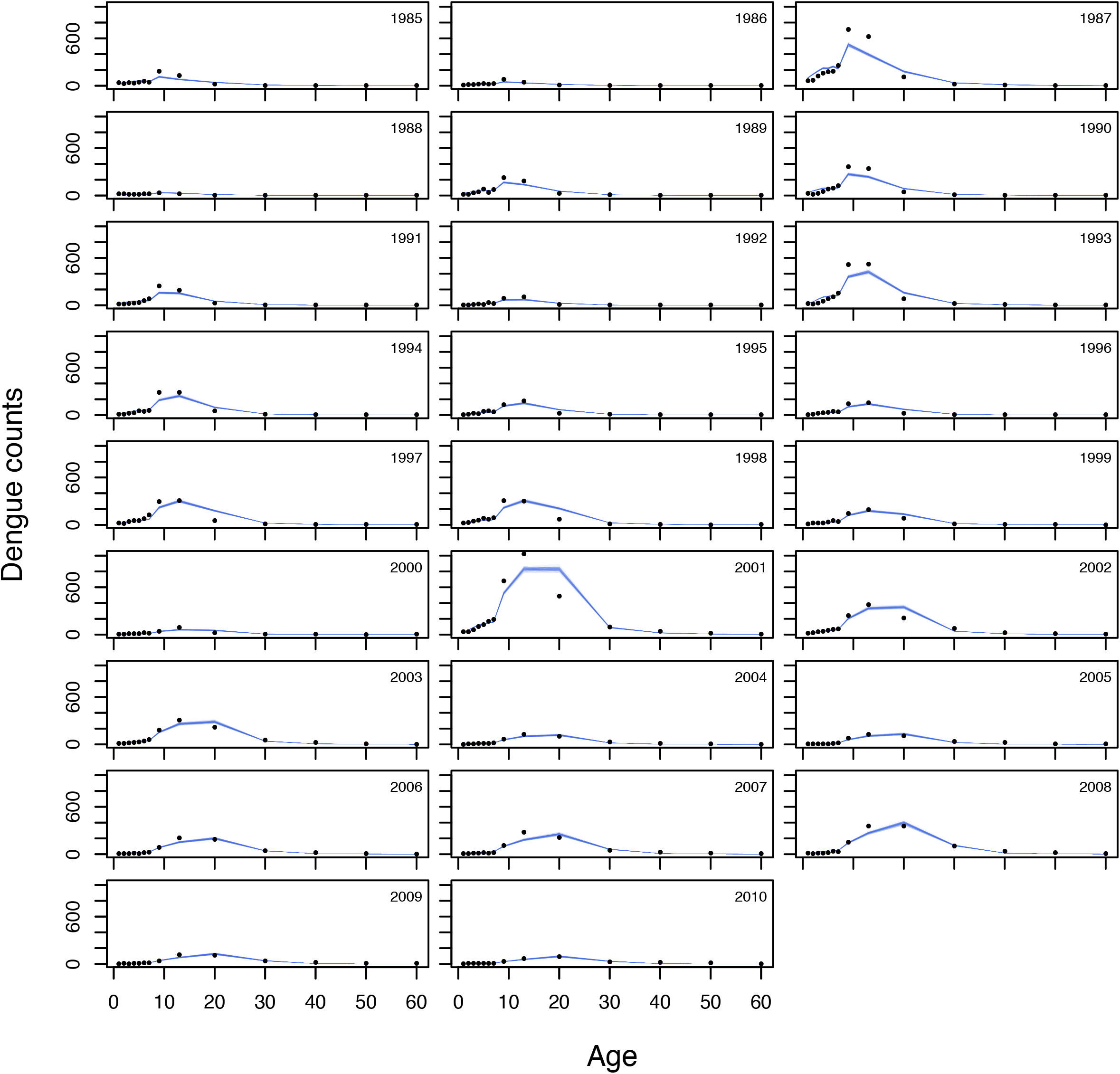

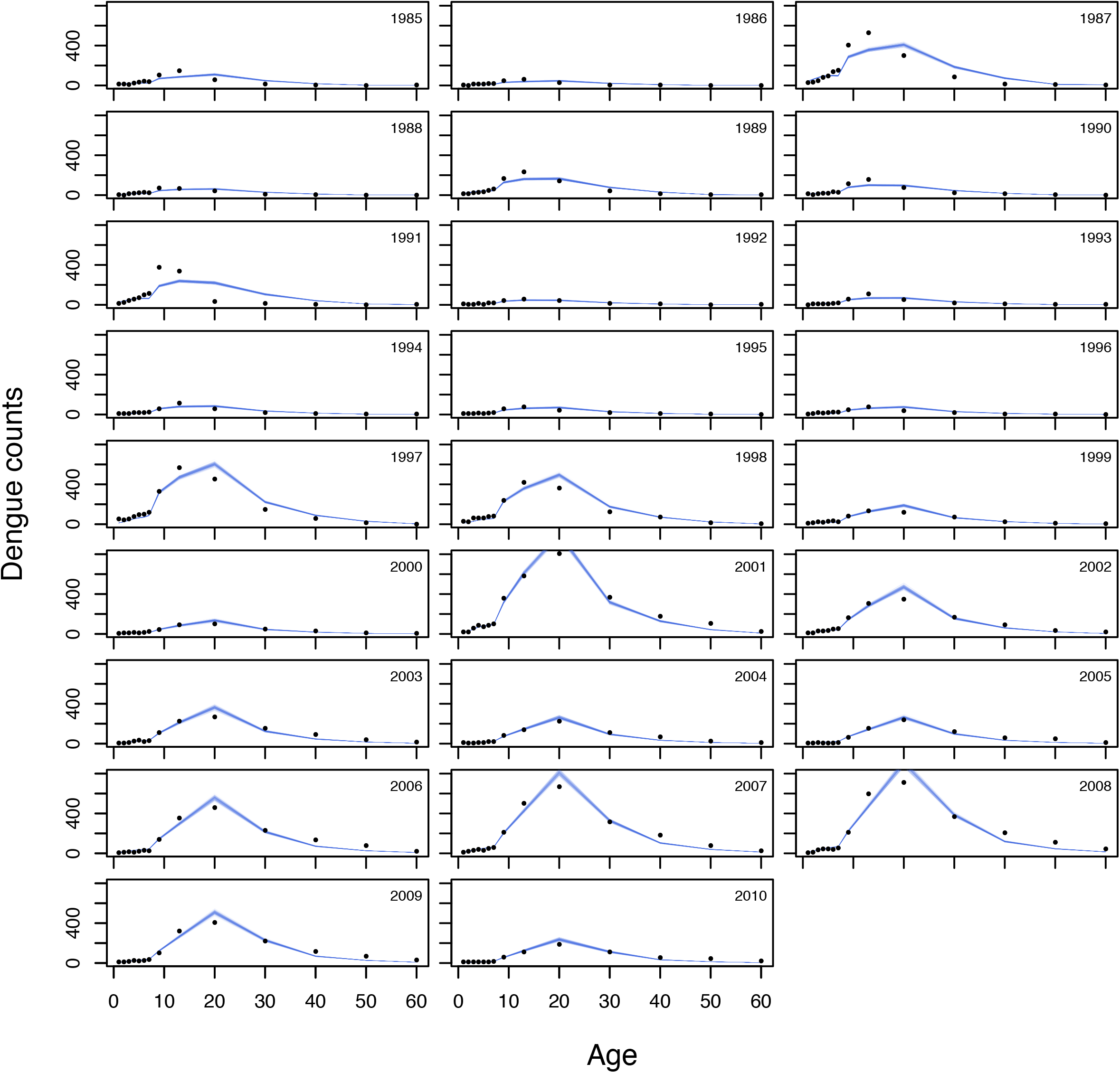

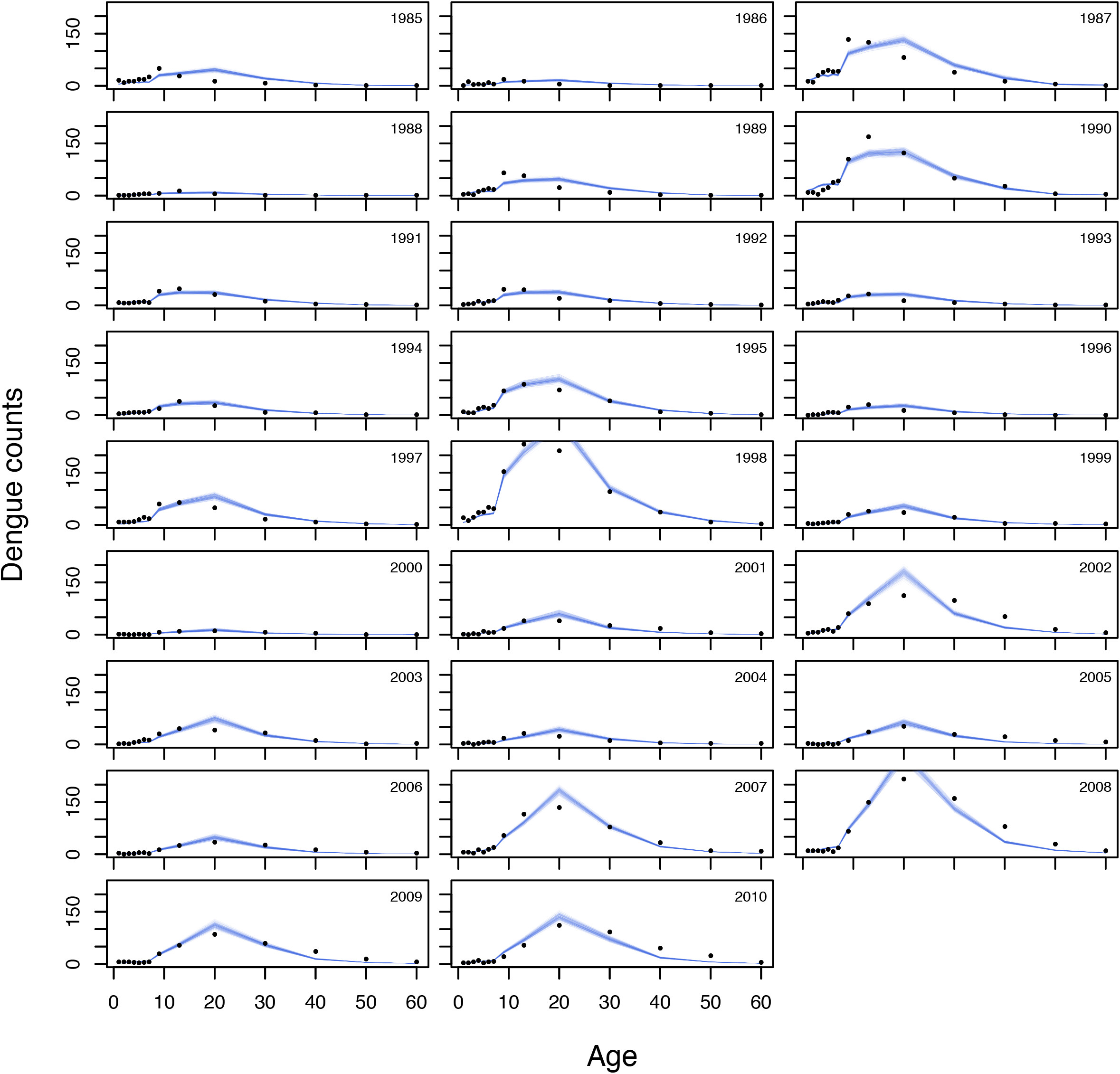

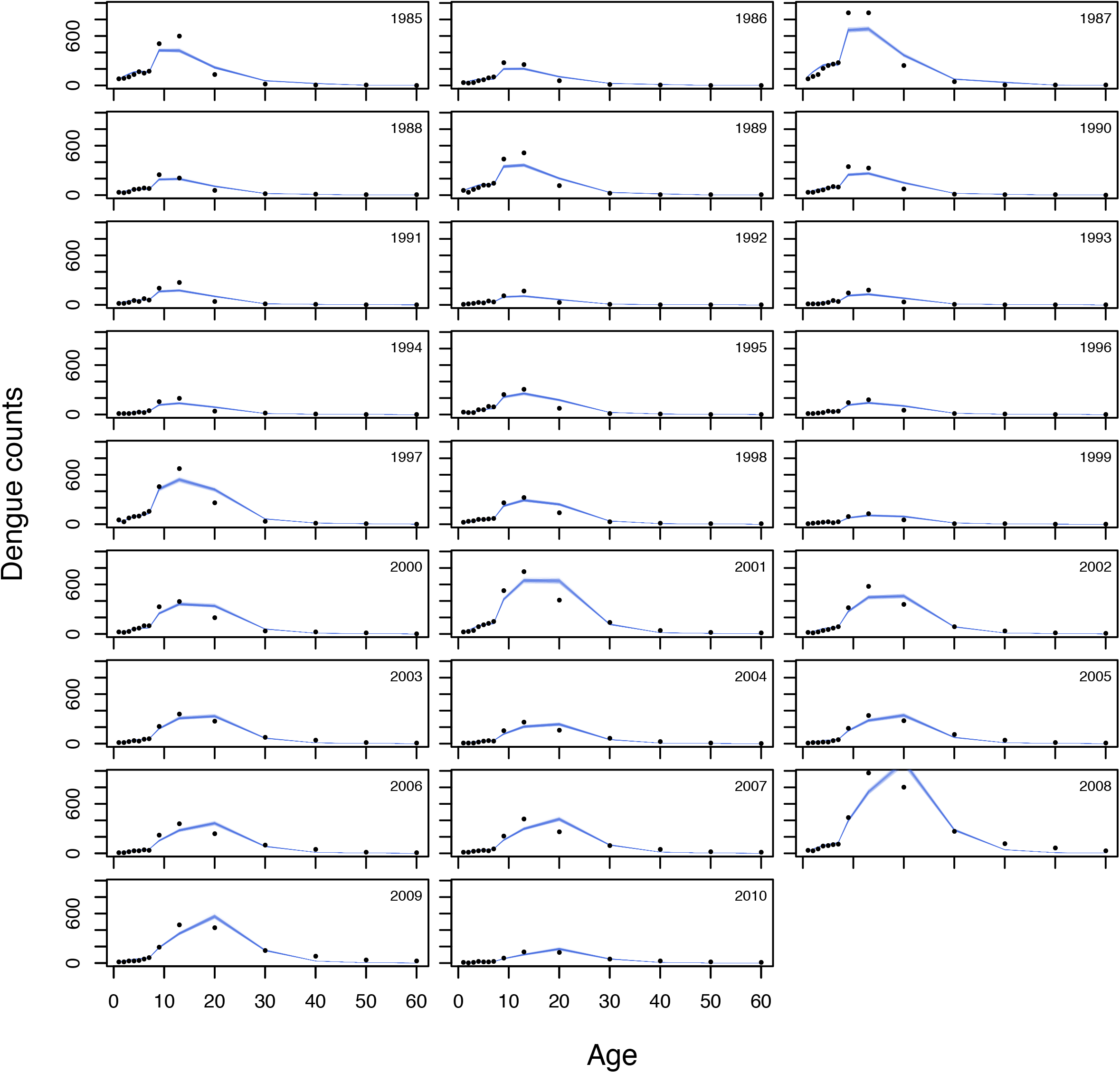

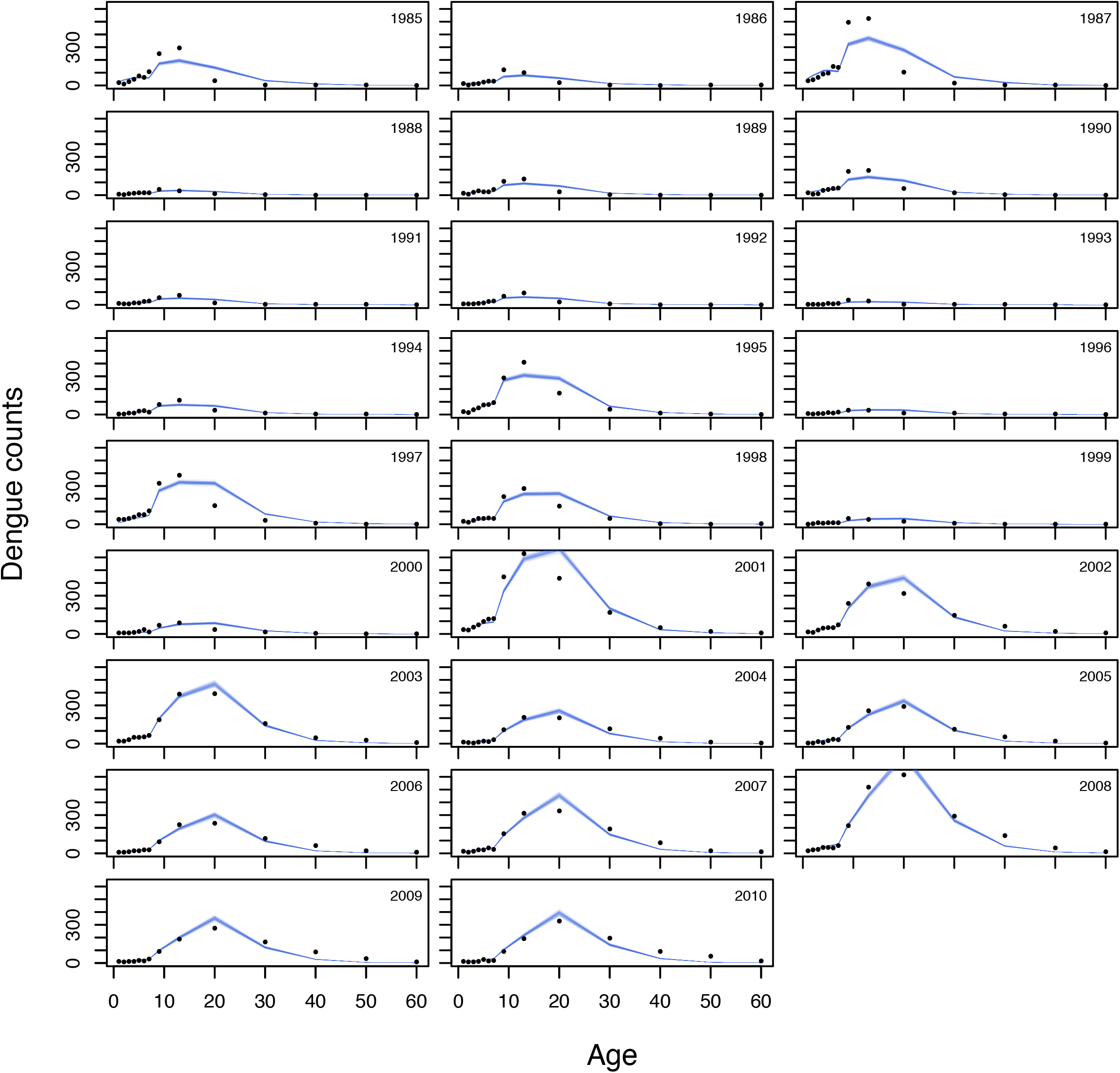

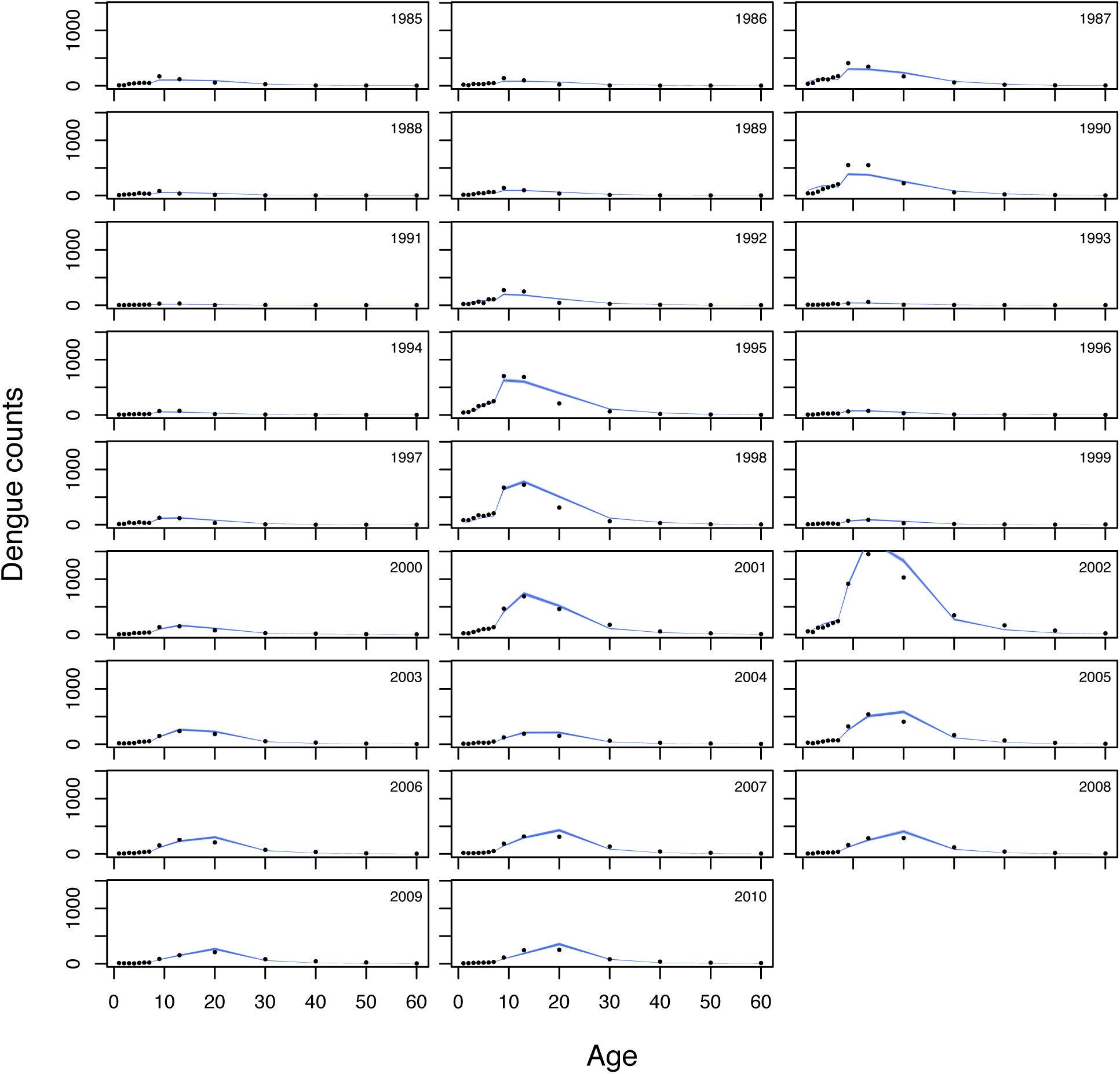

